# Asymmetric depth-filtration – a versatile and scalable approach for isolation and purification of extracellular vesicles

**DOI:** 10.1101/2021.08.23.457333

**Authors:** Vasiliy S. Chernyshev, Roman N. Chuprov-Netochin, Ekaterina Tsydenzhapova, Elena V. Svirshchevskaya, Rimma A. Poltavtseva, Anastasiia Merdalimova, Alexey Yashchenok, Amiran Keshelava, Konstantin Sorokin, Varlam Keshelava, Gennadiy T. Sukhikh, Dmitry Gorin, Sergey Leonov, Mikhail Skliar

## Abstract

A novel asymmetric depth filtration (DF) approach for isolation of extracellular vesicles (EVs) from biological fluids is presented, and its performance is compared with established methods. The developed workflow is simple, inexpensive, and relatively fast. Compared with ultracentrifugation and size-exclusion chromatography, the developed method isolates EVs with higher purity and yield. Only standard laboratory equipment is needed for its implementation, which makes it suitable for low-resource locations. The described implementation of the method is suitable for EV isolation from small biological samples in diagnostic and treatment guidance applications. Following the scale-up routes adopted in the biomanufacturing of therapeutics, which routinely rely on DF as one of the product purification steps, the developed method may be scaled up to harvesting therapeutic EVs from large volumes of growth medium.

## Introduction

Isolation of EVs from complex biofluids such as whole blood, plasma or serum, the growth medium of producer cells, urine, saliva, amniotic, and other biological liquids is challenging. EV isolations are often contaminated by a complex biomolecular milieu of biofluids and particles overlapping with EVs in size and other biophysical properties, such as lipid nanoparticle and protein agglomerates^1–4^. The yield of EVs isolated from such fluids varies significantly by isolation methods. For example, the yield ranges between 10^7^ and 10^13^ isolated EVs per mL of blood^2,5,6^. The purity of isolated EVs assessed by the protein concentration is similarly variable and was reported to be between 10^7^ and 10^11^ EVs per microgram of protein depending on the isolation method^7–13^.

Different isolation protocols lie on a tradeoff spectrum between unbiased isolation of all EVs from a biofluid and the purity of isolated EVs. On one extreme, precipitation techniques^14^ effectively pull all biological nanoparticles out of biofluid but at the cost of low purity and contamination by co-precipitated non-EVs content. On the other end of the spectrum, immuno-affinity capture^15^ isolates the least contaminated EVs but pulls only a minority subpopulation of vesicles that express the capture biomarker^16^. While the sample’s purity is essential, biased and fractionated isolations, even when pure, are problematic and may not accurately represent EVs’ biological activity, which may be multifactorial and multifaceted when the entire unbiased and heterogeneous population of EVs is considered^2,16^. Therefore, the isolation methods, including ultracentrifugation (UC) in its traditional^17,18^ and differential^19,20^ implementations with additional purification steps,^20–22^ size-exclusion chromatography,^23^ filtration,^24–26^ field-flow fractionation,^27^ and the combinations of such methods,^19,20,24,28^ may be assessed by their ability to reconcile the difficult task of maximizing the EV yield while minimizing cross contaminations^29^.

Filtration is a widely used separation technique generally divided into two categories: surface and depth filtration. Surface filtration strains particles that are too large to translocate through the filter while solubilized components and small particles pass through the pores. Larger particles accumulate on the upstream surface, forming a “cake” that eventually blocks the pore and impedes further filtration^30^. Surface filtration has already been used to isolate and concentrate EVs and remove such contaminants from the sample as microvesicles, macromolecules, and aggregates^7,26,31–36^.

The porous medium of depth filtration (DF) has pores too large to confine particles entirely on the filter’s surface. Instead, particles are fractionated kinetically based on the difference in their mobility through the medium. Solubilized components are eluted freely, while the transport of smaller particles is impeded but less than larger particles. The carrier flow transports small particles deeper into the filtration medium and elutes them first. Larger particles are either eluted at later times or trapped within the depth of the filter. Such trapping may be due to pores’ tortuous geometry, decreasing cross-section in asymmetric filters^38^, or the reduction in pores’ aperture by the already immobilized particles^30,39^. The accumulation of trapped particles eventually clogs the filter. However, since the entire medium participates in the straining, not just its surface, the depth filters can process much larger volumes of biofluid before clogging. Furthermore, the filtering capacity may be completely or partially regenerated by resuspending and eluting trapped particles by reversing the flow direction^30,40^. Overall, DF is an adaptable and scalable separation method with broad applications, including wastewater treatment^41–43^ and purification of manufactured biologics^44–47^.

Here, we apply the DF to isolate plasma EVs and demonstrate that its performance compares favorably with two established methods of EVs isolation - ultracentrifugation (UC) and size-exclusion chromatography (SEC). The developed method immobilizes EVs on the surface and within the depth of porous medium and then recovers them by reversing the carrier flow through the filter. To our knowledge, this is the first time the DF was implemented to isolate EVs. The method is direct, unbiased, and isolates high-purity EVs from complex biological fluids. It will find applications in liquid biopsy when EVs must be isolated from small volumes of biofluid and scaled up to process large volumes of growth medium during biomanufacturing of therapeutic EVs.

## Materials and methods

### Plasma sample

Human plasma was obtained with informed consent from a healthy adult at the Hospital of Pushchino Scientific Center of the Russian Academy of Sciences (Moscow Region, Russia), aliquoted into 1.5 mL Eppendorf tubes, and stored at –20° C until use.

### EV isolation by depth filtration

The DF is illustrated in Figure 1. Unlike surface filtration, the pores of depth filters at the flow’s entry surface (which we refer to as Surface 1) are larger than the particles we wish to exclude from elution. Instead, particles move through pores at a rate controlled by their size and surface interactions. Some depth filters have no distinct particle cut-off size. Smaller and less-interacting particles are eluted first, but all particles elute eventually if a carrier flow is maintained sufficiently long. In asymmetric filters, tortoise or narrowing pores trap larger particles within the filtration medium, but small particles can exit the membrane (Figure 1c).

**Figure 1:**
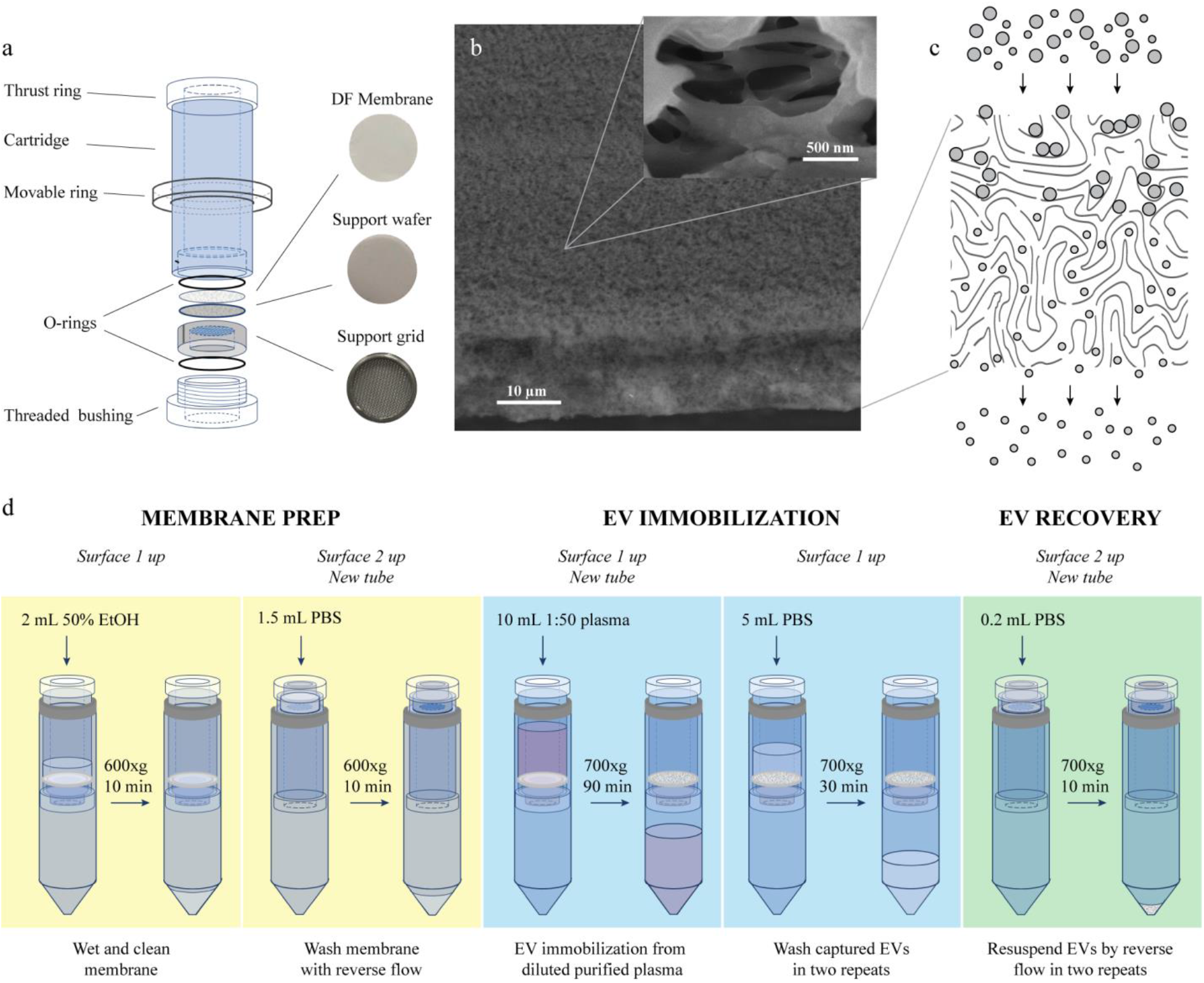
Depth-filtration cartridge and the protocol for DF isolation of EVs from blood plasma. (a) Cartridge components. (b) SEM image of the depth-filtration membrane showing its edge and Surface 1. Higher magnification (inset) of inlet pores on Surface 1 shows their aperture that is much larger than the size of EVs. As a result, the flow can drag vesicles inside the pores until they become immobilized within the depth of the filter. (c) Illustration of the depth filtration process showing two populations of particles of different sizes. Larger particles are retained within the volume of the filtration medium, while smaller particles are eluted. (d) Summary of the depth-filtration protocol used to isolate EV from blood plasma.

The membrane described here implements the latter type of DF, in which EVs are immobilized on the surface and within the depth of the filter, while proteins and other solubilized components of plasma elute with the carrier flow. EVs accumulate inside the filter and are later recovered by reversing the direction of centrifugally-driven flow by flipping the orientation of the filter assembly.

SEM image of Surface 1 and the cross-section of an asymmetric cellulose acetate DF membrane (∼20 μm thick), which we used to isolate EVs by depth filtration, is shown in Figure 1b. The asymmetry of the filtration medium is apparent from the SEM image of Surface 2 of the filter (Figure S8d in Supplemental Information). A disk of filtration medium, 22-mm in diameter, is held inside a cylindrical cartridge (19/25 mm ID/OD) fabricated from an acrylic resin (Prostagnost LLC (Patent # PCT/RU2017/000976), Moscow, Russian Federation) and designed to fit inside a standard 50 mL centrifuge tube (Figure 1a). The pore anisotropy of the selected DF membrane immobilizes EVs when the sample enters the filter through Surface 1 (forward flow direction) but then allows their recovery by reverse flow created by inverting the orientation of the cartridge relative to centrifugal forces.

The complete DF assembly (Figures 1a and S1 in SI) includes a metal sleeve with a stainless-steel grid supporting a porous wafer, which serves as a substrate for the DF membrane. A threaded plastic bushing is used to tighten the membrane securely in place. Two silicone O-rings on either side of the membrane are used to cushion the assembly from centrifugal forces (700-800xg) and prevent the fluid from bypassing the membrane during centrifugation. A movable stop ring keeps the assembly inside a centrifuge tube. It can slide along the cartridge between the thrust ring and threaded bushing. By changing the position of the stop ring and reversing the orientation in which the assembly is held inside a 50 ml tube, we invert the flow direction through the DF membrane.

Before proceeding to EV isolation, the DF membrane was wetted and flushed of potential contaminants. To that end, 2 mL of 50% ethanol was pipetted on the top surface of the membrane (Surface 1) and forced to flow through it by centrifuging at 600xg for 10 min (Figure 1d) or until EtOH completely passed through the filter. The cartridge was then flipped to reverse the membrane orientation (Surface 2 and small compartment of the cartridge facing up, Figure S8d), and the residual ethanol was driven out of the membrane by flowing 1.5 mL of 1x PBS by 600xg centrifugation for 10 min or until the entire volume of PBS passes through the filter (Figure 1d).

Blood plasma (200 µL for each repeat) was diluted 1:50 with PBS and centrifuged at 4,500xg for 30 min at 4° C to remove large impurities. The supernatant was then cleared by filtration through a 0.8 μm-pore cellulose acetate filter (Nalgene syringe filter, Thermo Scientific, Waltham, Massachusetts, United States), and 10 mL per DF cartridge were loaded on Surface 1 of the depth filter retained inside DF cartridge (Figure 1d). The cartridges containing aliquots were inserted into 50 mL tubes and centrifuged 70-90 min at 700xg using a swinging-bucket rotor until the sample has passed through the filter. At this point, the EVs were immobilized within and on the surface of the filter.

The immobilized EVs were washed by the direct flow of PBS to remove plasma background constituents, such as proteins, agglomerates, and lipid particles. For this purpose, 5 mL of PBS was flown through the membrane by centrifuging the cartridge at 700xg for 30 minutes. This step was repeated so that a combined 10 mL of PBS was forced through the filter to wash captured EVs.

The immobilized EVs were resuspended by reversing the flow through the membrane. The cartridges were flipped to change the filter orientation and inserted into new 50 mL tubes. The resuspending flow was created by centrifuging the assembly for approximately 10 min at 700xg to drive 200 μL of 1x PBS through the membrane in the reverse direction. This step was repeated to liberate additional EVs trapped inside the filter, which gave us ∼400 μL of the solution containing the isolated EVs for each aliquot of diluted plasma. This solution was then pipetted on and off Surface 1 of the filter to recover EVs remaining on Surface 1. Bubbles, often introduced by repeated pipetting, were removed by transferring the samples into 1.5 mL protein LoBind (Eppendorf) tubes and centrifuging for 15 min at 14,000xg. Degassed samples were then stored at -20° C until further analysis. We repeated the described DF isolation five times to test the reproducibility of EV yields.

### EV isolation by ultracentrifugation

Thirty mL of plasma was diluted 1:5 in PBS and aliquoted into five equal volume samples. The diluted fluid was transferred into 50 mL tubes and spun at 4,500xg at 4 °C for 30 min to pellet platelets, residual cells, and debris. The supernatant was transferred to new tubes, and the microvesicles were pelleted by 12,000xg centrifugation for 45 min at 4° C. The obtained supernatant was carefully transferred to 26 mL polycarbonate bottles, and small-sized EVs were isolated in two steps by ultracentrifugation using a 70Ti rotor (Beckman Coulter, USA). First, the samples maintained at 4° C were ultracentrifuged for 70 min at 100,000xg. The supernatant was discarded, and pellets resuspended in PBS in new 26 mL tubes. The second ultracentrifugation (100,000xg for 70 min) produced pellets, which we resuspended in 1 mL of PBS in 1.5 mL protein LoBind tubes and stored at -20° C until further analysis.

### EV isolation by size-exclusion chromatography

The EV isolation followed the protocol provided by the column manufacturer (PURE-EVs, HansaBioMed, Estonia). Briefly, the lower Luer slip cap was removed, and the column was washed with 15 mL PBS flowing at ∼1 mL/min. The lower Luer slip cap was then reinstalled, and remaining PBS above the column was removed. One mL of thawed plasma maintained at 4 °C was centrifuged for 30 min at 4,500xg, and 500 μL of supernatant was loaded into the prepared column. Luer-slip was removed, and the 30-second eluent fractions were collected. As effluent left the column, PBS was added to keep an uninterrupted flow. The flow rate stayed constant throughout the procedure at ∼1 mL/min, indicating the expected SEC operation. Isolation was repeated five times using different columns. Fractions enriched in EVs were pooled and stored at -20 °C.

### SEM imaging of immunolabeled EVs

The identity of particles in EV isolations was assessed by the expression of membrane proteins commonly associated with EVs (CD9, CD63, and EpCAM) and cell debris (Calnexin). Primary Abs – murine anti-CD63 (BioLegend, 353013), murine anti-CD9 (BioLegend, 312102), rabbit anti-EpCAM (Abcam, ab223582) and murine anti-Calnexin (LifeSpan, LS-B6014) – were diluted in PBS plus 0.5% bovine serum albumin (PBS-BSA; pH = 7.2-7.4) to 1:200 ratio. Dilutions of different Abs were separately mixed with EVs samples, and incubated for 14 hours at 4 °C. After the incubation, the samples were further diluted 1:5 in PBS-BSA, and unreacted antibodies were removed by centrifugal filtration (6,500xg) through a filter with ∼10-nm pores (∼100 kDa molecular weight cutoff). The Ab-labeled EVs retained by the filter were resuspended.

We used 20-nm gold nanoparticles to visualize CD9, CD63, EpCAM, and Calnexin expression in SEM images. Gold nanoparticles were functionalized with secondary antibodies – mouse or rabbit class G immunoglobulins (Abcam, ab27242 and ab27237) – designed to react with primary Abs, which we used to label EVs. We diluted Au nanoparticle suspension 1:1000 in PBS-BSA solution, mixed 200 μL of diluted suspension with 50 μL of EV samples, each labeled with a different primary antibody, and incubated the mixture for 6 hours at 4 °C. Unreacted gold nanoparticles were removed by filtration through a 30 nm filter. EVs and EV-Au-nanoparticle complexes retained by the filter were resuspended in 50 µL of deionized water. A small drop of the suspension (∼0.5 μL) was dried at ambient conditions on a clean silicon wafer. The wafer was placed on the SEM (Tescan MAIA3) specimen stage, and the desiccated sample was imaged using an accelerating voltage of ∼10.0 kV and magnifications between 100,000x and 500,000x. In the obtained SEM images, gold nanoparticles report the biomarker expression as bright spots on the surface of EV membranes.

### Nanoparticle Tracking Analysis (NTA)

The previously frozen samples were thawed and diluted in PBS to obtain the EVs concertation in the range suggested by the manufacturer of the NTA instrument (Nanosight model NS-300 equipped with 45-mW 488 nm laser; Malvern, Salisbury, UK). The required dilutions were between 1:100 to 1:1,000, depending on the isolation method. Within 1 min of the dilution, a sample was injected into the test cell and illuminated by the laser. The light scattered by particles was video recorded for 60 seconds by a high sensitivity sCMOS camera (camera level set to 14) at 25 frames per second. Each video consisted of 1,498 frames. Approximately 30-50 particles were observed in the field of view during video capture, corresponding to the concentration of ∼4-8×10^8^ particles per milliliter. The recording was repeated five times for each sample.

The videos were analyzed by Nanosight software (version 3.2) to measure the concentration of EVs, their size distribution, the mode and mean sizes, and the standard deviations of the results. Minimum track length, maximum jump mode, and blur size were set to Auto during video analysis. The detection threshold was set to 4. The viscosity of PBS was assumed to be that of water at the measured temperature. The sample’s temperature was measured automatically by the NTA instrument and stayed within 23-24 °C throughout the nanoparticle tracking experiments. The viscosity of water at this temperature is nearly constant at ∼0.91 cP.

### Dynamic Light Scattering (DLS)

Thawed samples were diluted 1:1,000 in PBS, and 1 mL of the preparation was transferred to a low-volume disposable sizing cuvette. After 5-min thermal equilibration inside the DLS instrument (Zetasizer Nano ZS, Malvern Instruments, Malvern, UK), the size distribution and ζ-potential of vesicles were measured at a 173° scattering angle, as recommended by the manufacturer for particles in the 0.3– 10,000-nm size range. The sample’s viscosity was assumed to be that of water. The measurements were interpreted by setting the refractive index of the solution to 1.33 and 1.35 for EVs^48^. Samples were analyzed in 5 repeats, each consisting of 12 light scattering measurements. The scattering data were processed assuming a general-purpose model implemented in the Zetasizer software, which produced the estimate of EVs ζ-potential, size distribution, mean, and standard deviations.

### Western Blotting

#### Extracellular vesicles and calnexin

Samples were separated on SDS-PAGE gel (Bio-Rad, Hercules, CA, USA cat. #456-1103) and electro-transferred to nitrocellulose membranes (Bio-Rad, Hercules, CA, USA, cat. #1704158) using Trans-Blot Turbo System (Bio-Rad, Hercules, CA, USA, cat. # 17001917). Membranes were washed with PBS and incubated overnight at 4° C with blocking buffer (Thermo Scientific, Waltham, MA, cat. #37572) to eliminate nonspecific sites. CD9, CD63, EpCAM, and calnexin expressions were detected on separate membranes, which were incubated overnight at 4 °C with primary antibodies diluted 1:5,000 in blocking buffer (BioLegend Way, San Diego, CA, cat. #312102 and #353013; LSBio, Seattle, WA, cat #B6014; Abcam, Cambridge, MA, cat. #ab223582, respectively) while shaking gently. After four washes, 10-min each, with 0.05% PBS-Tween 20 (PBST) solution, membranes were incubated at room temperature for 2 h in PBST-0.1% BSA solution of peroxidase-labeled secondary antibodies (IMTEK, Moscow, Russia, cat. #P-SAR and #P-GAM Iss) diluted 1:5,000. The incubated blots were washed (four times with PBST and then again two times with PBS; each wash was 10-min long) and developed using the Clarity Western ECL substrate (Bio-Rad, Hercules, CA, USA, cat. #170-5060). Immunoreactive bands were visualized with ChemiDoc™ XRS Imaging System (Bio-Rad, Hercules, CA, USA cat. #1708070).

#### Human serum albumin

Gel electrophoresis was performed in 10% PAAG using a Bio-Rad electrophoresis system. The transfer of proteins to Trans-Blot Transfer Media (Bio-Rad) nitrocellulose membrane was carried out using SemiDry Transfer Cell device (Bio-Rad). The membrane was blocked with 5% milk powder, washed in Tris buffer three times, and stained for 1 hour with shaking by using mouse anti-human albumin (Hy Test, Russian Federation, Moscow) diluted 1:1,000. Human serum albumin (Sigma, USA) was used as a control. After incubation, the membrane was washed three times with Tris buffer and incubated for one hour with anti-mouse antibodies conjugated with horseradish peroxidase (Santa Cruz Biotechnology, USA). After washing the membrane, the proteins were developed with a DAB/NiCl_2_ solution. The images were acquired using Gel Doc EZ Imager (Bio-Rad).

### Protein characterization

#### UV-Vis absorbance

Nanodrop 2000c (ThermoFisher Scientific, Waltham, MA, USA) was used to perform basic protein measurements. The characterization was performed using 1.5 uL of an undiluted sample, and the protein abundance was quantified using Nanodrop software (Protein A280 method). Each measurement was repeated four times.

#### Flow-cytometry

The presence of EVs was confirmed by using antibodies to human CD9-PE, CD63-APC, and CD81-FITC (Miltenyi Biotec, Germany). EVs were incubated with antibodies for 60 min at 4 °C. Unreacted antibodies EVs were removed by centrifugation at 15,000 rpm for 20 min, and the labeled vesicles were analyzed by flow cytometry (MACSQuant Analyzer, Miltenyi Biotec, Germany).

#### BCA protein analysis

Protein quantification using Micro BCA protein assay (Pierce™ BCA Protein Assay Kit, Sigma Aldrich, Poole, UK) was carried out following the manufacturer’s instructions. Briefly, EVs samples were diluted in DI water (1:1 ratio), and 150 µL of the solution was incubated for 2 h at 35° C with the equal volume of the reagent. Absorbance was then measured at 562 nm using a ClarioStar plate reader (BMG Labtech, Germany).

### Raman spectroscopy

The analysis was performed using a Raman spectrometer (Horiba LabRam Evolution HR, Horiba Ltd., Irvine, California) equipped with Olympus M Plan 50X objective and 600 lines/mm grating. Raman scattering was excited by a 633-nm laser adjusted to 50% of its maximum power. A small drop (∼1 µL) of EV sample was pipetted on a fused quartz surface and dried at room temperature. The analyte concentration was increased by placing the second drop in the same location and drying. Three spectra were accumulated for averaging, each obtained with a 50-second exposure. Clean area of quartz glass and the dried solution of human serum albumin (0.4 g/mL; Octapharma Pharmazeutika Produktionsgesellschaft m.b.H., Austria) were used as controls.

### Data analysis

Size-frequency measurements obtained by different techniques were converted into probability density functions (pdf) and visualized as histograms.

## Results

### EV isolation by depth-filtration

The concentration of EVs isolated by DF varied between 2 and 3×10^11^ particles/mL in different aliquots (Figure 2b, Table S1). The vesicles’ hydrodynamic size distributions measured by NTA and DLS are shown in Figures 2a, S2, and S3. The mean hydrodynamic diameters for these distributions were 109 ± 2 nm and 97 ± 2 nm, respectively (Tables 1 and S2). The vesicles’ surface charge was characterized by their ζ-potential, which was –12.4 ± 0.5 mV.

**Table 1:**
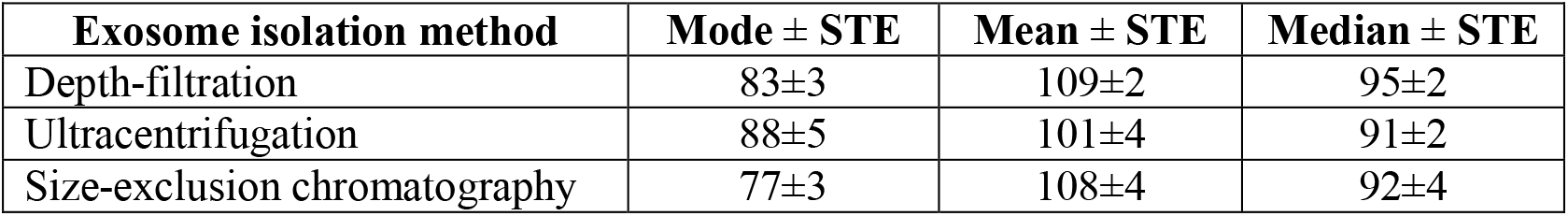
Mode, mean, and median hydrodynamic diameters (nm) of extracellular vesicles (plus-minus standard error, STE) of five repeated isolations by different methods.

**Figure 2.**
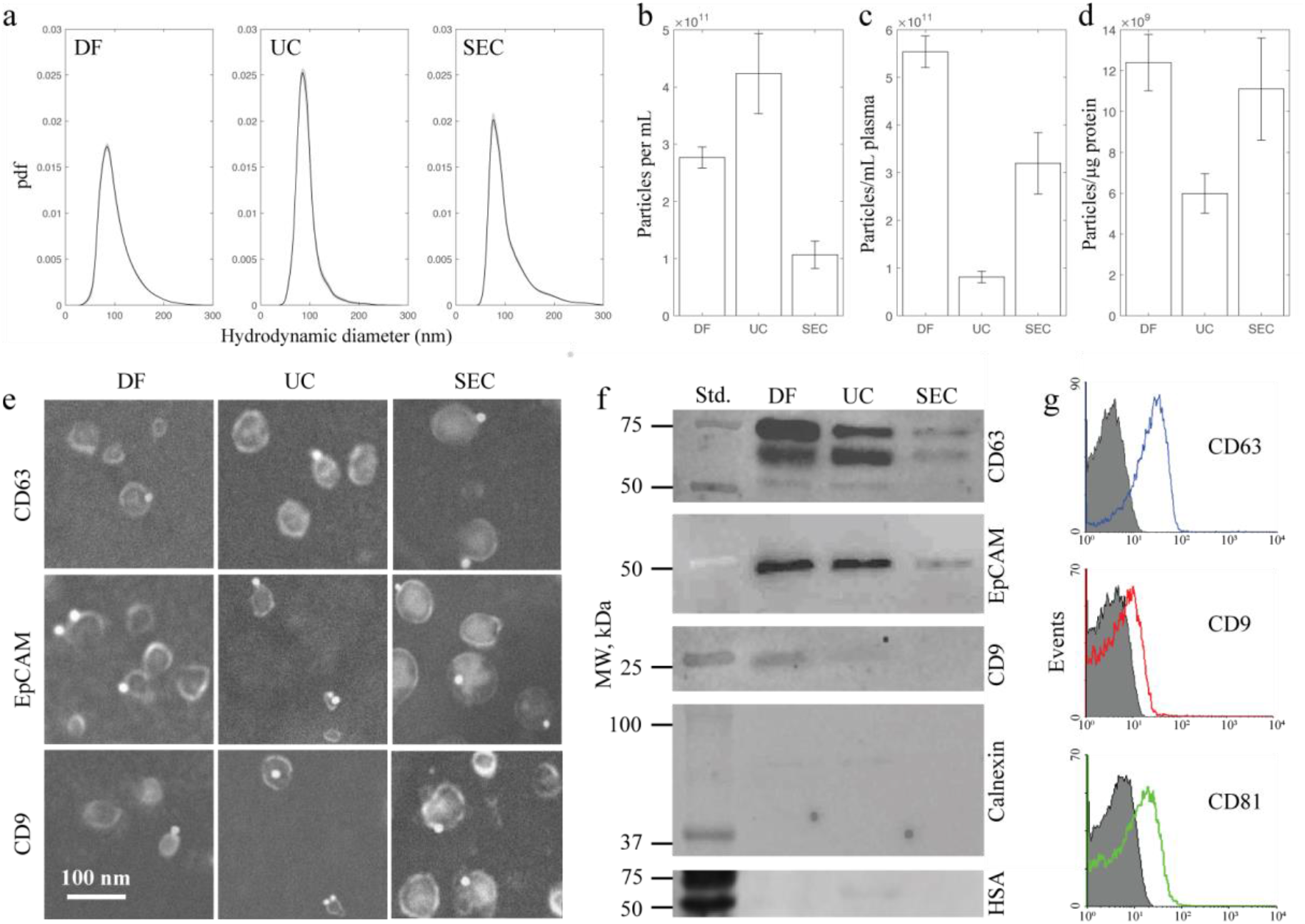
Characterizations of EVs isolated by DF, SEC, and UC. (a) The size distribution of plasma EVs isolated by different methods expressed as the probability distribution function (pdf). (b) The NTA measurements of EV concentration in different samples. (c) EVs isolated per mL of plasma. The yield of depth filtration is substantially higher compared to the alternatives. (d) The protein abundance per extracellular vesicle assesses the purity of EV isolations. The purity was comparable in DF and SEC isolations and lower in UC samples. (e) SEM images of EVs labeled with CD9, EpCAM, and CD63 primary antibodies. Bright dots on the EV membrane are 20-nm gold nanoparticles reporting the locations of biomarker expression. (f) Western blots show the highest expression of exosomal biomarkers CD63, EpCAM, and CD9 in the EV sample isolated by DF. Cross contaminations quantified by negative controls (calnexin and human serum albumin) are the lowest in SEC and DF-isolated samples. (g) Flow cytometry analysis of biomarkers expressed in DF-isolated EVs.

SEM was used to image the morphology and the size of EVs’ membranes. Figure 2e and S4 show the expected rounded shape of desiccated vesicles^48^ and the geometric size of membrane envelopes smaller than their hydrodynamic diameter^49^. The expression of CD9, CD63, and EpCAM biomarkers is reported in SEM images by immunolabeled gold nanoparticles, which appear in Figure 2e as bright spots on EV membranes.

BCA-measured protein concentration was between 21 and 29 μg/mL in the five experimental repeats, which corresponds to 1.1–1.4×10^10^ vesicles per microgram of proteins. EVs expressed canonical EV biomarkers, CD63, EpCAM, and, to a lesser extent, CD9, as was determined by Western blotting (Figure 2f) and flow cytometry (Figure 2g). The most abundant plasma protein, albumin, was undetectable (Figure 2f), indicating that DF isolation effectively eliminates solubilized plasma proteins from the isolated EVs. Calnexin, an endoplasmic reticulum integral protein, also was not found, suggesting that DF-isolated EVs were not contaminated by cell debris.

Raman spectrum of EVs isolated by DF (Figure S5) agrees with previous reports by Krafft et al.^50^ and Slyusarenko et al.^51^ and contains the expected peaks assigned in Table S3. Specifically, the peak near 704 cm^-1^ corresponds to cholesterol and cholesterol esters. The spectral region between 1,200 and 1,300 cm^-1^ corresponds to amide III bands, and the peak at 1,440 cm^-1^ is due to CH_2_ bending in lipids and cholesterol.

### EV isolation by UC

We followed the UC protocol described in references^52,53^. The particle sedimentation by centrifugal forces depends on the particle size, buoyant density, and viscosity of the solution. We reduced the sample’s viscosity by diluting plasma with PBS^54,55^, which improved the sedimentation efficiency. We pelleted and discarded cells, cell debris, apoptotic bodies, large microvesicles, and aggregates by a two-step conventional centrifugation at 4,500xg and then at 12,000xg^53^. EVs were isolated from the supernatant by ultracentrifugation. A visible pellet was saved and resuspended in PBS. The pelleting by UC was repeated to reduce the protein contamination. The second pellet was again resuspended in 1 mL of PBS and saved for analysis.

The yield of plasma EVs isolated by UC was almost an order of magnitude lower than by DF (Figure 2c). The distribution of hydrodynamic sizes (Figure 2a and Table 1) was not significantly different. The ζ-potential equal to -11.3± 0.8 mV was comparable to the electrokinetic potential of vesicles isolated by DF. The EV biomarkers (Figure 2f) were less expressed than after DF isolation, while the total protein concentration was higher. The purity of UC samples quantified as the number of vesicles per microgram of proteins was below 1×10^10^ particles/μg (Figure 2d). The Western blot in Figure 2f indicates that albumin is one source of a higher protein concentration^56^. Albumin was undetectable in DF- and SEC-isolated samples.

### EV isolation by SEC

Commercially available SEC columns were used to isolate EVs from 500 μL aliquots of plasma. As the sample flows through a gel-packed column containing porous resin beads, the propagation paths of particles are size-dependent. Smaller particles migrate through the pores that retard their translocation. Particles too large to enter pores migrate through the gel filling the volume unoccupied by beads and elute first. This mechanism separates particles of different sizes by their elution time.

We used NTA measurements to determine that particles with hydrodynamic diameters between 90-100 nm were predominantly eluted from the column during 30-second intervals number 7, 8, and 9, assuming that the first interval starts when 500 μL of plasma is loaded in the column (Figure S6). Later-eluting fractions contained smaller particles (mean size in the range between 50 and 60 nm) and higher protein concentrations quantified by Nanodrop measurements (A280 protein analysis, Figure S7). The concentration of co-isolated proteins in pooled fractions 7-9 measured by BCA protein assay was lower than in UC-isolated samples (Figure 2d). Fraction 10 contained EVs at a lower abundance relative to protein concentration (less than 1×10^10^ particles/μg of protein) and overlapped with the protein elution time (Figure S7). Therefore, fraction 10 was excluded.

EV concentration in pooled fractions 7-9 was in the range between 0.8×10^11^ and 1.4×10^11^ particles/mL (Figure 2b, Table S1), which is ∼3 times lower than the concentration obtained by DF. Mean particle size determined by DLS (115 ± 9 nm) and NTA (108 ± 4 nm) was consistent with DF and UC isolations. The ζ-potential was -10.8 ± 0.4 mV, close to the values obtained for EVs isolated by alternative methods. The protein concentration in pooled SEC fractions measured by BCA protein assay was 9.5±0.2 μg/mL. After the normalization to the concentration of isolated vesicles measured by NTA, the relative abundance of co-isolated proteins was low – a microgram of protein was found in a volume containing 1.1×10^10^ ± 2.5×10^9^ EVs – and only slightly higher than in DF isolation, Figure 2d. Non-EV proteins calnexin and albumin were not present in SEM and DF-isolated samples (Figure 2f).

## Discussion

We quantitatively compared the performance of DF with two commonly utilized methods – UC and SEC. The yield of EVs per mL of human plasma was significantly higher by depth filtration (Figure 2c). Therefore, out of the three examined isolation methods, the depth filtration is the least biased^8,14,57^, providing the most accurate representation of extracellular vesicles present in the blood. EV biomarkers (CD63, CD81, CD9, and EpCAM) were most expressed in DF-isolated samples. The contamination by plasma proteins and membrane debris, respectively indicated by HSA and calnexin expressions, was the lowest (Figure 2f).

We selected an asymmetric cellulose acetate membrane to perform EV isolation by depth filtration due to its unique characteristics. Cellulose acetate has a negative surface charge^58^, which impedes nonspecific adsorption of biological molecules with a negative ζ-potential. For this reason, cellulose acetate filters have been used for over 60 years in the purification and isolation of DNA, other nucleic acids, and proteins,^59,60^ including by DF^42^. In our case, the negative charge prevents the surface adsorption of EVs that have a negative ζ-potential at neutral acidity, as previously reported^61,62^ and confirmed in this study. Low surface adsorption of EVs contributes to the high yield of the DF isolation and low contamination by proteins, small non-membrane (e.g., lipid) particles, and membrane fragments. These contaminants, not bound by adsorption, are easily removed from the filtration medium by the direct flow during EV washing, after which the reverse flow resuspends the immobilized EVs.

We used two types of synthetic nanoparticles to gain insight into the isolation mechanism and the performance of the described DF method. The first sample was a suspension of 100-nm size-standard NIST-traceable polystyrene (latex) beads (Polysciences, Inc., Warrington, PA, USA; Figure S8b) diluted in PBS to 1×10^11^ particles/mL. Latex beads are known to have a negative ζ-potential, and their size was selected to be close to the mean diameter of isolated plasma EVs (Table 1). The suspending buffer, void of a complex molecular milieu of plasma, transits the DF filter with ease. We, therefore, modified the protocol for isolating latex beads from the PBS, as shown in Figure S8a. Specifically, lower 400xg forces and shorter (7 to 8 min) centrifugation were sufficient to drive the entire 3.75 mL sample through the filter. Only a small fraction of 100-nm particles (less than 1%) transited through the filter with the permeate (Figure S8c). Approximately 88% of the particles were recovered after 2 mL of PBS was repeatedly pipetted and aspirated off the filter’s Surface 1. The recovered particles were likely retained on the top surface of the membrane and accumulated close to pore entrances. Indeed, the SEM image of the filter (top image in Figure S8d), taken after the latex sample was flown through it, shows many particles present on Surface 1 and inside pores (inset in Figure S8d). No particles were observed on the exit Surface 2 nor inside the pores within its immediate proximity. The reverse flow of 2 mL PBS, created by 5-min centrifugation at 400xg and repeated three times, sequentially recovered 1.7, 0.3, and 0.1% of latex beads in the original sample. The remaining ∼10% of the particles were permanently lodged within the filter’s depth and could not be recovered.

The second synthetic sample was a suspension of 20-nm gold nanoparticles functionalized with anti-mouse IgG (ab27242, Abcam, Cambridge, MA, USA) diluted in PBS to 1×10^11^ particles/mL. These particles with protein-decorated surfaces were selected to test if smaller-than-EVs particles and solubilized proteins pass through the depth filter without contaminating particles in the EV range retained by the filter. Indeed, we found that Au particles easily transit through the filter. Approximately 90% of them eluted with the permeate (Figure S8c) when the protocol applied to the suspension of latex beads was followed. Very few Au particles were recovered by the reverse flow of PBS, which we repeated three times until 6 mL of resuspending fluid was flown through the filter.

In summary, this report describes a novel approach to EV isolation by depth filtration. The developed method is simple and inexpensive. It reproducibly isolates EVs from complex biological fluids in 3 to 4 hours using only basic laboratory equipment, such as a conventional centrifuge capable of producing 700xg centrifugal forces. Therefore, it may be used in point-of-care applications and even implemented with manually-powered centrifugation.^63^ The main components of the DF cartridge can be reused after cleaning, and only the DF cellulose acetate membrane must be replaced before each isolation. The method may be scaled up by simultaneously processing multiple centrifuge tubes up to the capacity of a laboratory rotor. Further scaled up is possible by specialized centrifugation or using a pressure-driven flow to harvest clinically meaningful quantities of therapeutic EVs from large volumes of growth medium.

## Authors’ contributions

V.S.C. and M.S. conceived the study, designed experiments, and wrote the manuscript. V.S.C. conducted EV isolation, tests with synthetic particles, NTA and SEM analysis. R.N.C-N. and E.T. performed Western-blot analysis and BCA. E.V.S. and R.A.P. conducted flow cytometry and Western-blot analysis of human serum albumin. A.M. and A.Y. performed Raman spectroscopy and peak identification. A.K., K.S. and V.K. assisted in the design and fabrication of the DF cartridge. G.T.S., S.L. and D.A.G. provided laboratory equipment and guidance. Everyone contributed to the manuscript revision.

## Acknowledgements

Authors would like to thank KDSI (Malvern Panalytical Ltd representative in the Russian Federation) for providing the instrument for nanoparticle tracking analysis (Nanosight NS300). Authors also would like to acknowledge the Hospital of Pushchino Scientific Center of the Russian Academy of Sciences for providing plasma with volume sufficient to conduct this study.

## Funding

This work was supported by the Russian Science Foundation under Grant № 20-73-00102.

## Declaration of interest statement

The authors declare no competing interests.

## Notes

### Competing Interest Statement

The authors have declared no competing interest.

