## Supplemental material for "Asymmetric depth-filtration – a versatile and scalable approach for isolation and purification of extracellular vesicles"

### Supplementary material

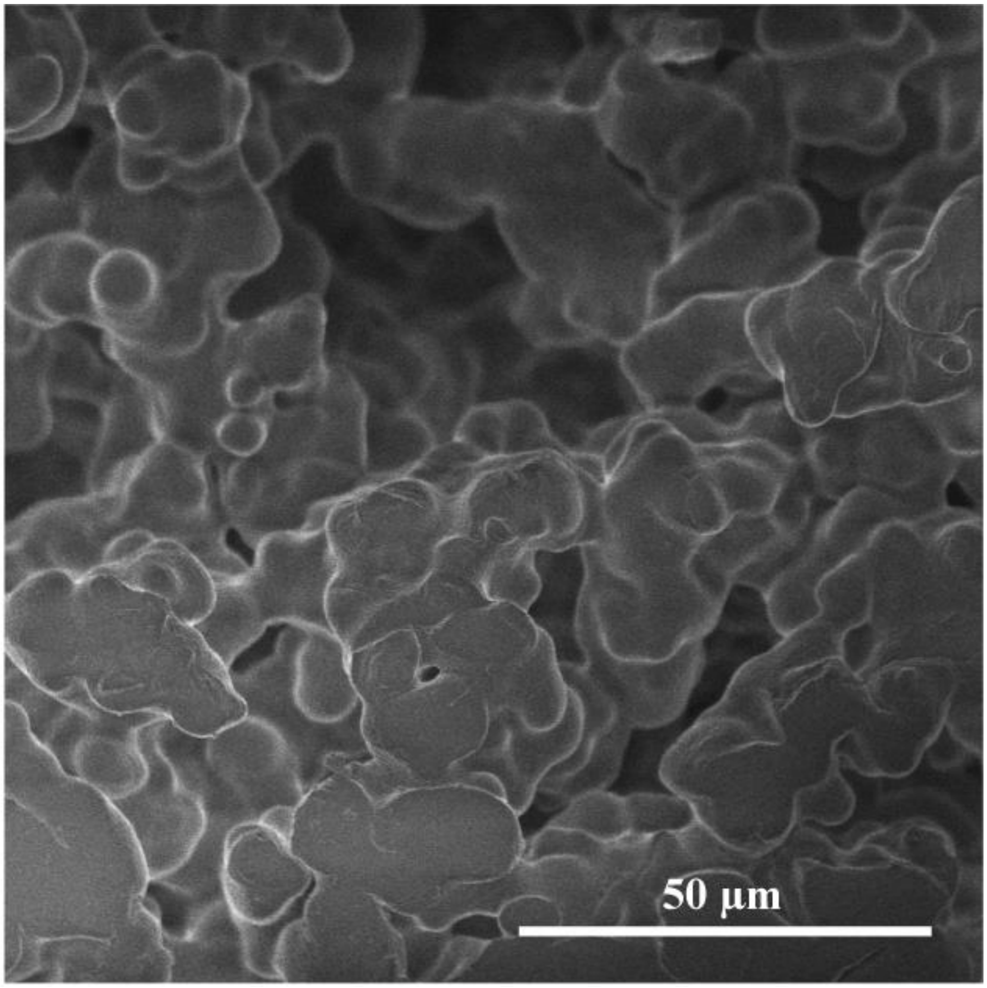


**Figure S1.** Scanning electron microscopy image of the base wafer used to support the cellulose acetate DF membrane.

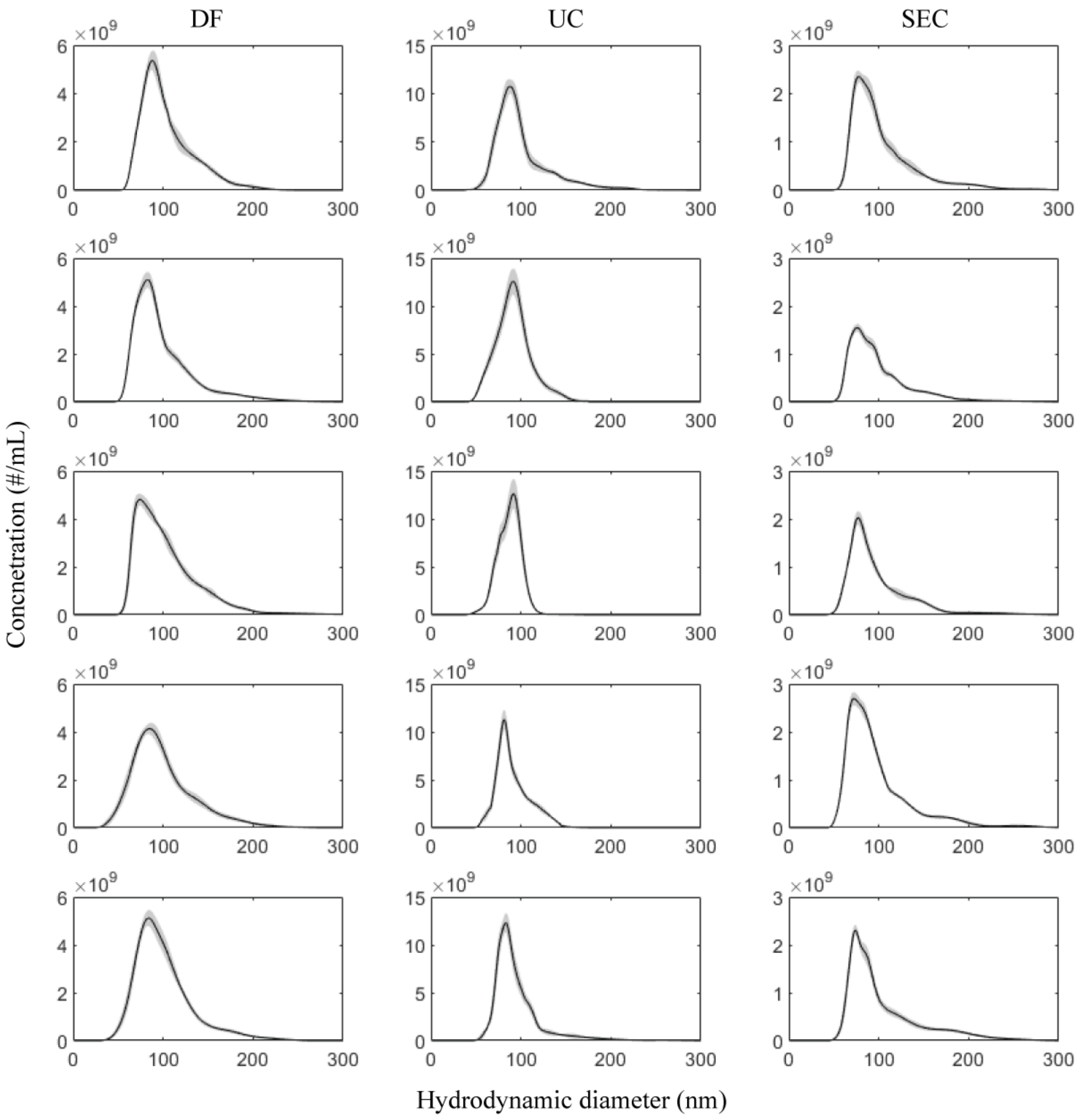


**Figure S2.** NTA measurements of hydrodynamic size distribution and concentration of EVs isolated by different methods.

**Table S1.** NTA measurements of EV concentration after the isolation by different methods (#/mL).

| Repeats | DF | UC | SEC |
|---|---|---|---|
| Run 1 | 2.90E+11 ± 1.80E+10 | 3.79E+11 ± 9.30E+09 | 8.95E+10 ± 3.08E+09 |
| Run 2 | 2.72E+11 ± 1.79E+10 | 3.39E+11 ± 9.10E+09 | 1.40E+11 ± 2.61E+09 |
| Run 3 | 3.01E+11 ± 3.16E+10 | 4.16E+11 ± 1.30E+10 | 1.07E+11 ± 3.47E+09 |
| Run 4 | 2.64E+11 ± 4.06E+10 | 4.70E+11 ± 6.85E+09 | 1.18E+11 ± 1.06E+10 |
| Run 5 | 2.56E+11 ± 3.85E+10 | 5.14E+11 ± 1.05E+10 | 7.90E+10 ± 5.51E+09 |

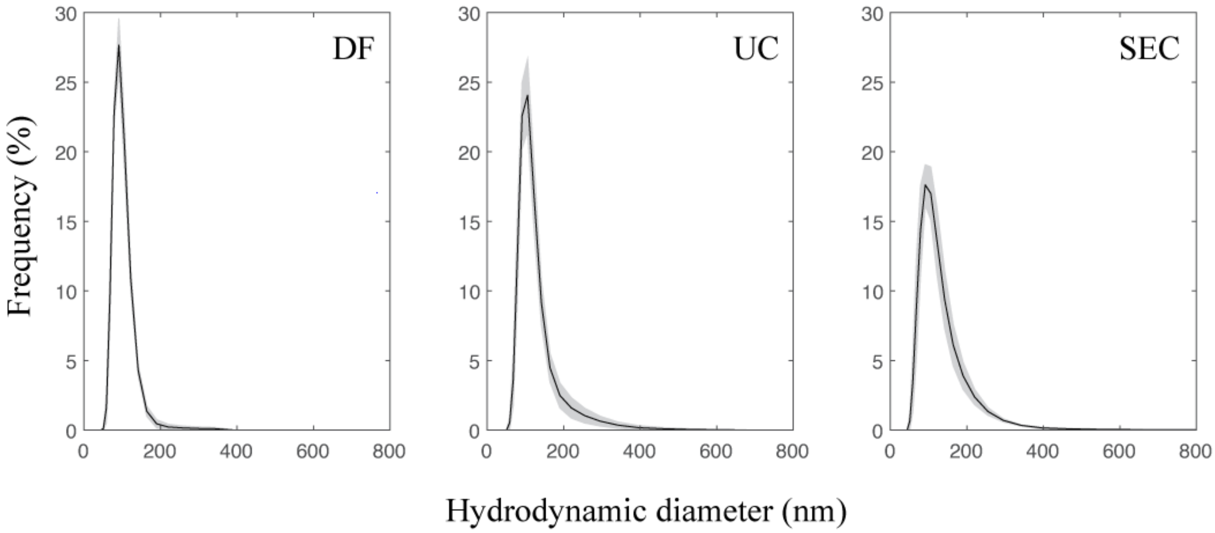


**Figure S3.** Hydrodynamic size distribution of isolated EVs determined by DLS. The solid lines show the mean value after five repeats. The standard deviation is shown by shading.

**Table S2.** Mean hydrodynamic diameter (five repeats) and ζ-potential (three repeats) plus-minus the standard deviation (STE) of isolated EVs determined by DLS.

| Repeats | Mean hydrodynamic diameter ± STE (nm) | ζ-potential (mV) |
|---|---|---|
| Depth-filtration | 97 ± 1 | -12.4 ± 0.5 |
| Ultracentrifugation | 115 ± 6 | -11.3 ± 0.8 |
| Size-exclusion chromatography | 115 ± 9 | -10.8 ± 0.4 |

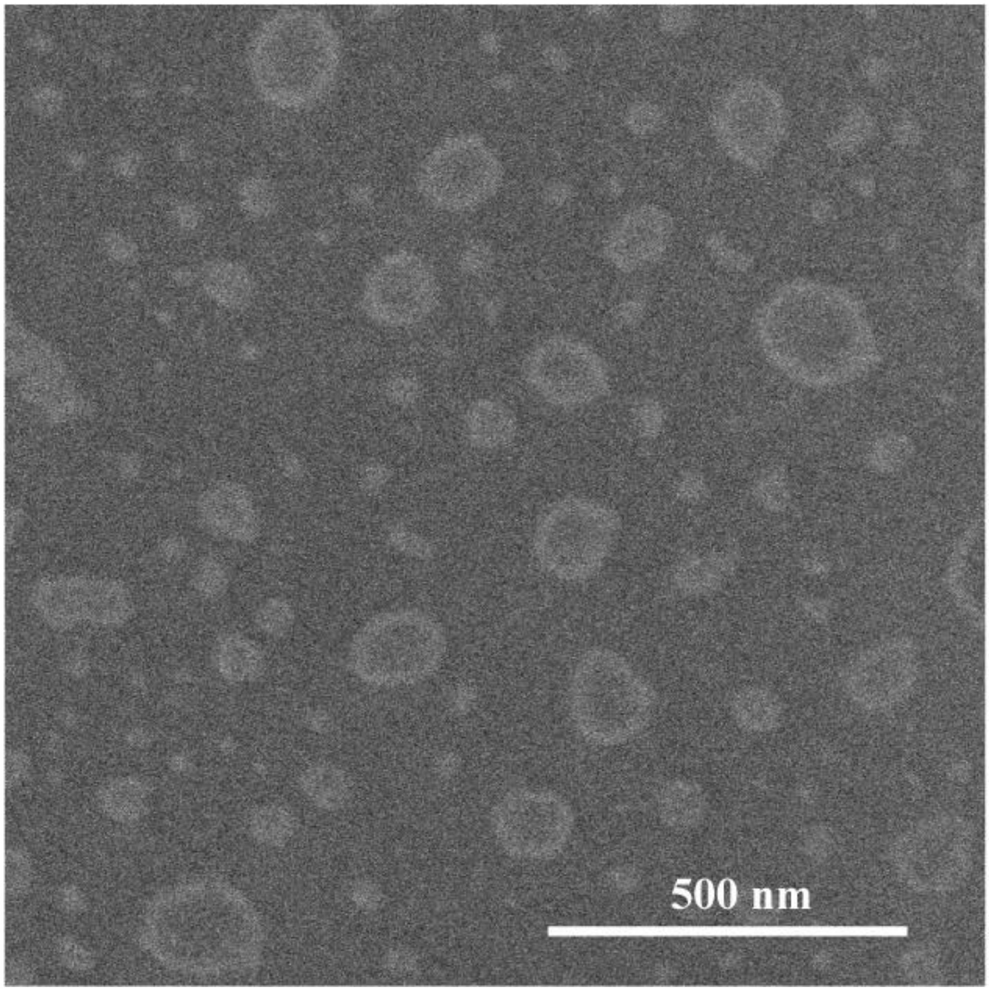


**Figure S4**. SEM image of desiccated EVs isolated by asymmetric depth filtration.

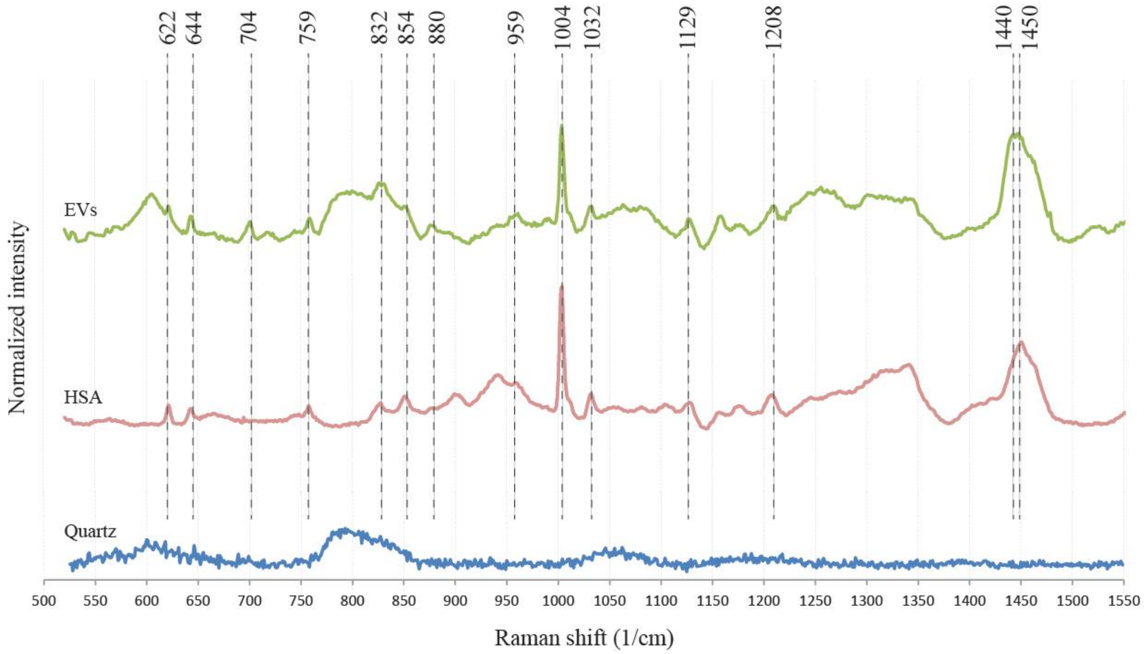


**Figure S5**. Raman spectrum of EVs isolated by depth-filtration is compared with reference spectra of human serum albumin (HSA) and fused quartz substrate. The interpretation of observed peaks is given in Table S3.

**Table S3.** Raman peaks identified in EV spectrum.

| Raman shift, $cm^{-1}$ | Assumed assignment | Reference |
|---|---|---|
| 622 | Phenylalanine (phenyl ring breathing) | [1] |
| 644 | Tyrosine (C–C twisting) | [2] |
| 704 | Cholesterol and cholesterol esters | [3,4] |
| 759 | Tryptophan | [5] |
| 832 | Tyrosine (out of plane ring breathing) | [2,5] |
| 854 | Tyrosine (ring breathing mode); proline (C-C ring stretch) | [5] |
| 880 | Tryptophan; in-plane rocking ($CH_2$), e.g., protein | [2,6] |
| 959 | Cholesterol | [2,5] |
| 1004 | Phenylalanine, CC aromatic ring stretching | [1,3] |
| 1032 | $CH_2CH_3$ bending (e.g., phospholipid); C-C vibration (e.g., polysaccharide) | [6] |
| 1129 | Lipids and proteins | [3] |
| 1208 | Phenylalanine, tryptophan (C-$C_6H_5$ stretching) | [7]; [5] |
| 1200-1300 | Amide III in proteins | [3] |
| 1440-1450 | Lipids $CH_2$ deformation at 1437; lipids and proteins $CH_2/CH_3$ deformation at 1443; protein $CH_2$ bending mode at 1446 | [2]; [5]; [6] |

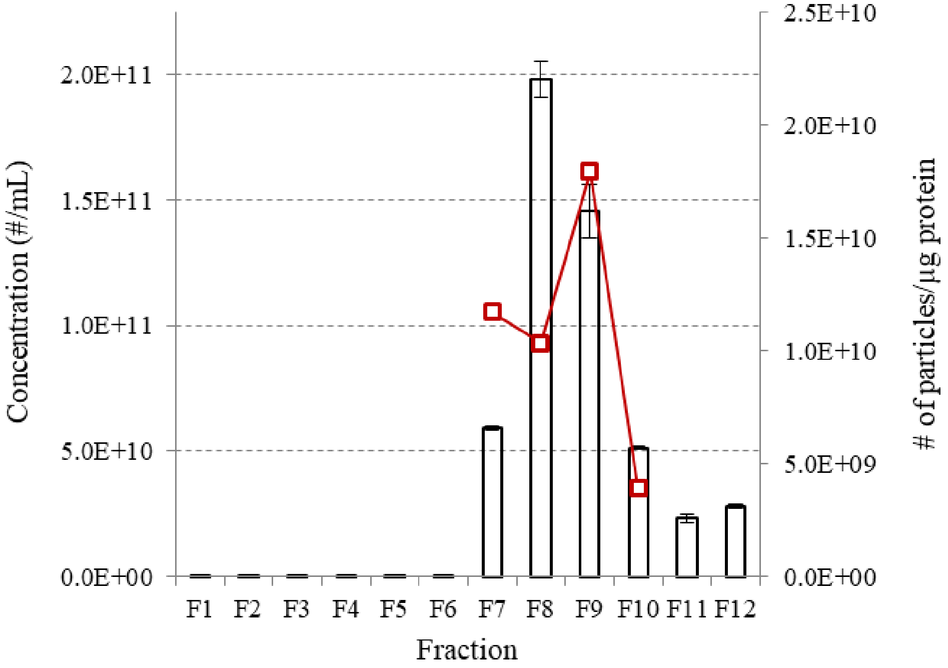


**Figure S6.** NTA and BCA results for different 30-second SEC fractions. Fractions 7-9 have particle concentrations near 1E10 particles/µg protein (red graph, right scale). Fractions 7-9 were pooled for subsequent analysis and compared with EV isolations by depth-filtration and ultracentrifugation. The shown particle concentration is adjusted for dilution needed to perform NTA measurements.

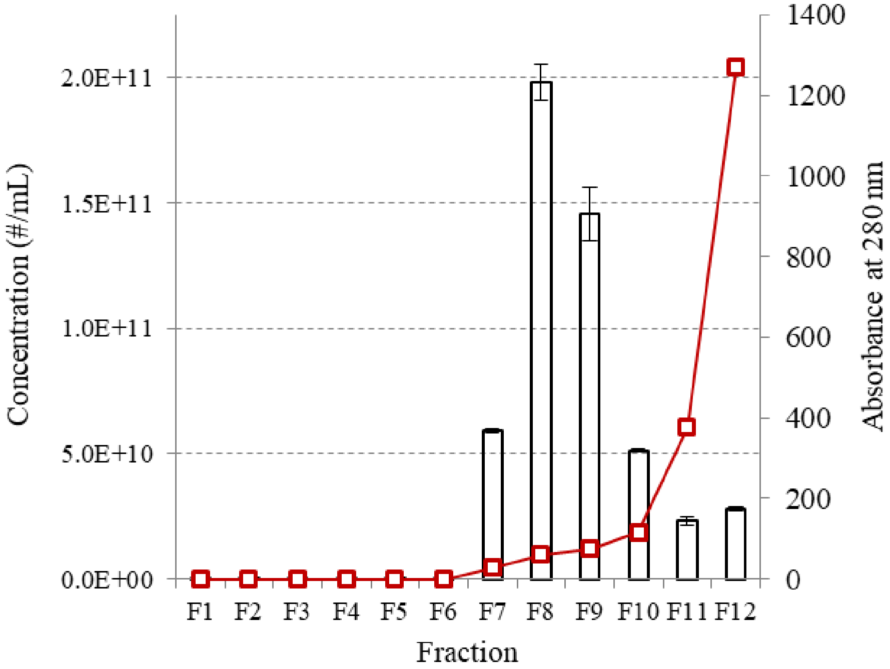


**Figure S7.** Results of NTA and Nanodrop Protein A280 measurements for SEC fractions. Fractions 7-9 have the optimal tradeoff between particle and protein concentrations (red graph, right scale). A steep increase in absorbance starting with fraction 10 indicates a rapid rise in protein contamination.

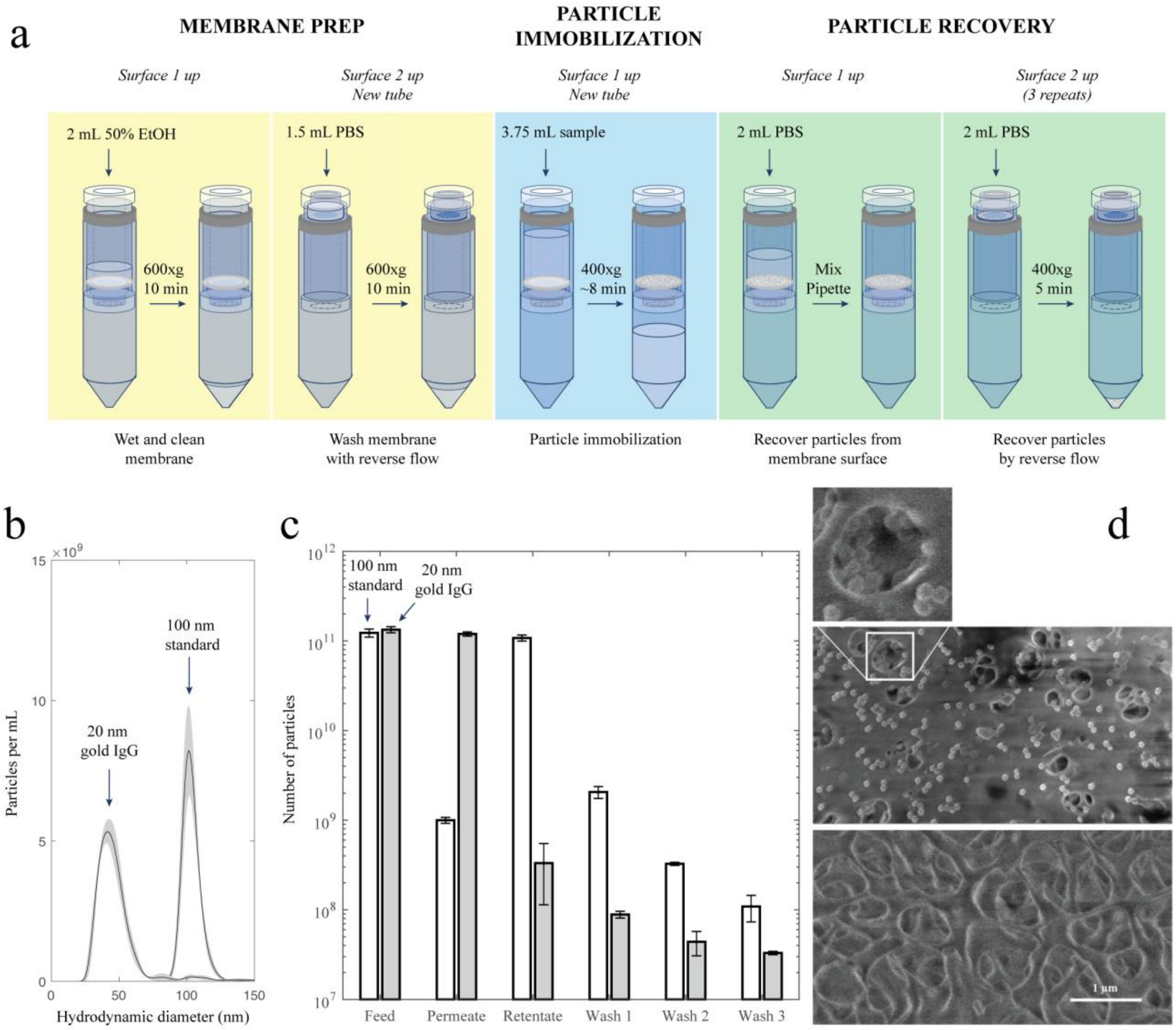


**Figure S8.** Depth-filtration of synthetic samples: 100 nm nanobeads and 20 nm gold nanoparticles functionalized with anti-mouse IgG. (a) The depth-filtration protocol was modified for synthetic samples which transit through the membrane with less resistance than plasma samples. (b) Hydrodynamic diameters of latex standards and Au particles. (c) Particle concentration in the original synthetic samples (Feed) is compared with the number of particles that passed through the filter, retained on its top surface (Surface), and resuspended by the reverse flow applied sequentially three times. (d) SEM imaged Surface 1 of the cellulose acetate membrane (top image) after 100-nm standard beads were flown through it. Latex beads are seen on the filter's entry surface and inside the pores (details in inset). The bottom image shows Surface 2 of the membrane.
